# SpliceVec: distributed feature representations for splice junction prediction

**DOI:** 10.1101/183087

**Authors:** Aparajita Dutta, Tushar Dubey, Kusum Kumari Singh, Ashish Anand

## Abstract

Identification of intron boundaries, called splice junctions, is an important part of delineating gene structure and functions. This also provides valuable insights into the role of alternative splicing in increasing functional diversity of genes. Identification of splice junctions through RNA-seq is by mapping short reads to the reference genome which is prone to errors due to random sequence matches. This encourages identification of splicing junctions through computational methods based on machine learning. Existing models are dependent on feature extraction and selection for capturing splicing signals lying in vicinity of splice junctions. But such manually extracted features are not exhaustive. We introduce distributed feature representation, *SpliceVec*, to avoid explicit and biased feature extraction generally adopted for such tasks. SpliceVec is based on two widely used distributed representation models in natural language processing. Learned feature representation in form of SpliceVec is fed to multi-layer perceptron for splice junction classification task. An intrinsic evaluation of SpliceVec indicates that it is able to group true and false sites distinctly. Our study on optimal context to be considered for feature extraction indicates inclusion of entire intronic sequence to be better than flanking upstream and downstream region around splice junctions. Further, SpliceVec is invariant to canonical and non-canonical splice junction detection. The proposed model is consistent in its performance even with reduced dataset and class-imbalanced dataset. SpliceVec is computationally efficient and can be trained with user defined data as well.

## 1. Introduction

In most of the eukaryotes, a protein coding gene is interrupted by intervening sequences called introns. These introns are removed from the exonic sequences via splicing process, which occurs cotranscriptionally [1]. The process generates mature substrates to be translated into proteins. Splicing involves snipping of introns at exon-intron and intron-exon junctions, called *splice sites* or *splice junctions*, and ligation of exons. Identification of splice junctions is therefore an important part of delineating gene structure and functions. Gene identification as well as its structural annotation has become an important problem in bioinformatics due to the abundance of sequenced genomes made available by advanced sequencing technologies like RNA-seq. RNA-seq, in recent times, has also provided meaningful insights into the role of alternative splicing (AS) in increasing functional diversity of genes. AS is a regulated process which alternatively skips or joins exons, or parts of exons or introns to form variety of proteins during translation. Recent estimates by RNA-seq suggest that more than 90% of multi-exon genes in human body undergo alternative splicing [2], thus making the identification of splice junctions all the more crucial. Identification of splice junctions from RNA-seq involves mapping of millions of short reads to the reference genome. However, multiple potential match of the short read on the reference genome makes sequence mapping less reliable [3]. Also, the existing alignment based methods [4, 5] consider detection of splice sites based on only canonical splicing patterns (GT at donor site and AG at acceptor site) thereby missing important non-canonical splicing patterns [6].

The vast availability of annotated sequences makes it possible to create large enough training datasets for supervised learning algorithms to predict splice sites. The correct prediction of splice sites is facilitated by the identification of relationships and dependencies among the nucleotides around the splice sites. This is motivated by the observation that the splicing signals are most likely to reside in the vicinity of splice sites [7]. The learning algorithms need a set of features for training the model. Creating an optimized set of features that best represent the dataset has always remained a challenge for splice site prediction. The presence or absence of certain nucleotide sequences close to the splice sites were considered as features for splice site prediction for a long time [8, 9, 10, 11, 12, 13, 14]. Since all such features were not known, there have been constant efforts to improve or refine features as well as include more relevant features by taking into account recent experimental observations. The large number of features may not only contain many irrelevant features but may also adversely affect classifier performance due to its high dimension. This led to many efforts for obtaining the optimum feature set through feature selection [15, 16, 17, 18]. But the set of features obtained were not exhaustive.

This scenario motivated the adoption of an approach that can represent features specific to splice junctions based on the splicing signals without any manual extraction and selection of features. Rather, the model itself will capture motifs from the biological sequences that act as splicing signals for splice site selection. Lee et. al. proposed a deep Boltzmann machine based methodology for splice junction prediction [6]. Recently, Zhang et. al. have employed a deep convolutional neural network (CNN), named *DeepSplice* [19], that learns features that characterize the true and decoy splice junctions. They have predicted novel splice junctions based on these features and obtained state-of-the-art performance of 96% accuracy. A distributed representation of biological sequences was proposed by Asgari et. al. [20] and Kimothi et. al. [21] for protein family classification task where a prediction accuracy of more than 99% was achieved.

This paper introduces a novel approach for distributed feature representation of splice junctions by embedding it in an n-dimensional feature space. Each dimension in the feature space represents one feature of the corresponding splice junction. This embedding, named *SpliceVec*, is in the form of n-dimensional continuous distributed vector representation. The embeddings are learned by a shallow neural network using unsupervised data. We explored two variants of SpliceVec, namely *genome based SpliceVec (SpliceVec-g)* and *splicing-context based SpliceVec (SpliceVec-sp)* for feature representation of splice junctions. We evaluate the quality of SpliceVec in both intrinsic and extrinsic tasks. For the intrinsic evaluation, we visually inspect two dimensional representation of true and false splice sites using Stochastic Neighbor Embedding (t-SNE) [22]. We evaluate SpliceVec on splice junction classification task for the extrinsic evaluation. In contrast to the recent deep learning methods, we use simple multilayer perceptron (MLP) as a classifier. We name this model as *SpliceVec-MLP*.

Our results and contributions can be summarized as follows:

- We propose two variations (SpliceVecg and SpliceVec-sp) for feature representation of splice sites. SpliceVec outperforms state-of-the-art methods by 2.42-18.86% in terms of accuracy for splice site prediction.
- We explored the optimal sequence length that best captures the splicing signals for improving the prediction results. We find that inclusion of entire intronic sequence significantly boosts the predictive power of the classifier.
- The proposed feature representations are more robust in handling reduced training samples. SpliceVec maintains an accuracy above 99% even with a 60% reduction of training samples whereas the accuracy of its counterpart drops by about 6%.
- SpliceVec is more consistent in its performance with class-imbalanced data making it more suitable for the real time scenario where number of pseudo sites are several times more than that of true splice sites.
- SpliceVec-MLP identified non-canonical splice junctions (junctions not comprising of the consensus dimer GT or AG at donor or acceptor site respectively) with 100% accuracy indicating that our feature representations are invariant to both canonical and non-canonical splice junctions.
- SpliceVec-MLP can be deployed in both CPU and GPU environment. SpliceVec-MLP, being 12.94 times computationally faster than the state-of-the-art model, contributes as a suitable option for classification of the abundant annotated sequences available these days by high-throughput sequencing technologies.

## 2. Methods

The proposed approach can be divided into two stages: the feature representation stage that generates a distributed representation for each splice junction based on either of the two frameworks, namely *word2vec* and *doc2vec*, and classification of splice junctions using MLP. We shall discuss all these in the following subsections. An overview of the proposed approach is shown in Figure 1.

**Figure 1:**
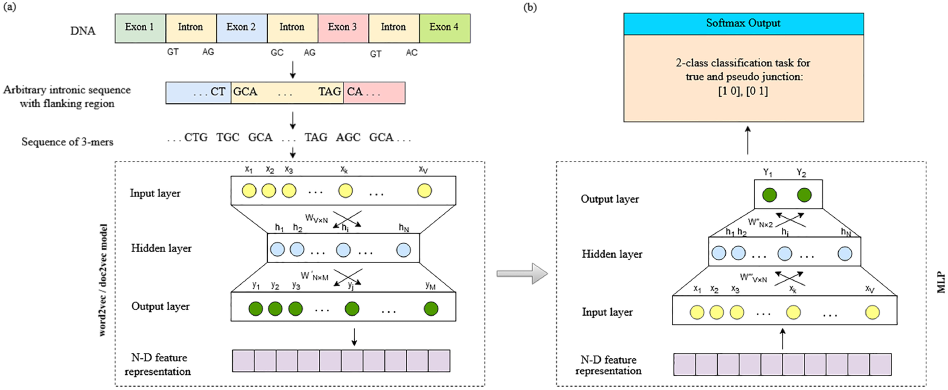
Proposed approach. (a) feature representation; (b) splice junction classification.

### 2.1. Data

We have used the latest release of GENCODE annotation [23] (version 26), based on human genome assembly version *GRCh38*, for extracting true and false splice junctions. This version was released in March 2017. We extracted 294,576 splice junctions from the protein coding genes. Based on these splice junctions, we observed that an intron length varies from 1 to as much as 1,240,200 nucleotides (nt). A recent study suggests that the shortest known eukaryotic intron length is 30 base pairs (bp) belonging to the human *MST1L* gene [24]. Introns shorter than 30bp are usually accounted to sequencing errors in genomes. Based on this study, we considered only those introns whose length were greater than 30 bp. This reduced the number of splice junctions to 293,889. We selected 293,889 false splice junctions by randomly searching for splice site consensus sequences GT and AG which were not annotated as splice junctions. This was considered as a necessary condition for selection of false splice junctions because more than 98% of splice junctions are canonical, that is, they contain the consensus dinucleotide GT at the donor site and AG at the acceptor site [25]. In DeepSplice, the length of false splice junctions was considered to be lying between 10 and 300,000 nt, the reason of which was not clear. So, instead, we considered only those false splice junctions whose length was not less than 30 nt and not more than 1,240,200 nt with both donor and acceptor splice sites lying in the same chromosome. We considered this range to mimic the scenario of true introns.

### 2.2. Distributed representation

Vector representation of a word or a sentence is an integral part of many natural language processing (NLP) tasks. Although local representations, like N-grams and bag-of-words, have been successfully applied in NLP, they lack efficiency due to sparsity and high dimensionality. Most importantly, such representation needs to be defined explicitly. In the recent times, distributed representation has been most successful in complex NLP tasks. This type of representation is learned based on the connection and interaction between words appearing in various contexts within a chosen corpus.

With this idea, Mikolov et. al. proposed word2vec models [26] that compute continuous vector representations for words learned by shallow neural networks. These models embed each word in an n-dimensional space where syntactically and semantically similar words appear close to each other. The model has two different architectures-continuous bag of words (CBOW) and skip-gram. CBOW predicts the current word given some surrounding context words whereas skip-grams predicts context words given the current word.

This model has been further extended in the form of doc2vec model [27] to incorporate continuous vector representations for variable length texts like sentences, paragraphs and documents. This model also comes in two architectures-distributed bag of words (DBOW) and distributed memory (DM). DBOW, similar to skip-gram architecture of word2vec model, predicts the context words from the document vector. DM, on the other hand, predicts the current word based on surrounding context words as well as the document vector. This is similar to CBOW architecture of word2vec model. In doc2vec model, the word vectors are global and shared among documents whereas the document vectors are local to the document and learned only in context of the corresponding document.

### 2.3. Distributed representation of splice junctions

Both the models, described in the previous section, demand as input a large corpus of text and produce outputs in n-dimensional vector space where each unique word in the corpus is assigned a vector in that space. A corpus in NLP is a continuous chain of words following certain grammatical structure. On the other hand, biological sequences are a continuous string of four characters, *A, C, G*, and *T*, representing nucleotide bases Adenine, Cytosine, Guanine and Thymine respectively. Our corpus also contains *N* which can represent any of the four nucleotides. Since there is no concept of words in case of biological sequences, it can be broken down into k-mers of any length k. There have been experiments with variable length k-mers [28] as well as both overlapping and non-overlapping k-mers [20, 21] for different bioinformatics problems. In this paper, we focus on overlapping 3-mers. For example, a biological sequence ATTGGCA will yield the following sequence of words: ATT, TTG, TGG, GGC, and GCA. Thus, our vocabulary can have a maximum of 5^3^ = 125 distinct words. After this pre-processing step of fragmenting biological sequences into words, we fed the corpus into both word2vec and doc2vec models to generate n-dimensional embedding, named SpliceVec. SpliceVec-g is based on word2vec model whereas SpliceVec-sp is based on doc2vec model.

#### 2.3.1. Genome based SpliceVec

This type of feature representation is generated using word2vec model. For word2vec model, our corpus is the complete human genome assembly (version *GRCh38.p10*). We broke down the genome assembly into chromosomes. We have considered 24 chromosomes, that is, chr1 to chr22 as well as X and Y chromosomes. We have excluded the mitochondrial and unlocalized sequences because of its exceptionally small size. We further fragmented each chromosome into sentences of 2000 characters. Each sentence was broken into overlapping 3-mers representing words. We trained word2vec model with the complete sequence of 3-mers. The word2vec model produced an n-dimensional vector representation for each unique 3-mer in the corpus. We next computed the vector representation of a splice junction by taking the average of summation of the vector representation of each word in the splice junction sequence. We used gensim [29] library of python to generate the word vectors.

#### 2.3.2. Splicing-context based SpliceVec

This variation of feature representation is based on doc2vec model. For doc2vec model, our corpus is the complete set of true and false splice junctions. Each splice junction, considered as a document, was broken down into overlapping 3-mers to form words and fed into doc2vec model as training data. The model generated a vector representation for each unique word and each splice junction sequence in an n-dimensional hyperspace. We used the C-implementation of doc2vec model by Le and Mikolov [27] for generating the vectors.

Both the CBOW and skip-gram architecture for word2vec model, as well as the DBOW and DM architecture for doc2vec model are illustrated in Figure 2 considering an arbitrary biological sequence ATTGGCA. The sequence is broken down into 3-mers sequence ATT, TTG, TGG, GGC, and GCA where TGG is the current word and remaining four words are context. ID refers to the splice junction which contains the given sequence of 3-mers.

**Figure 2:**
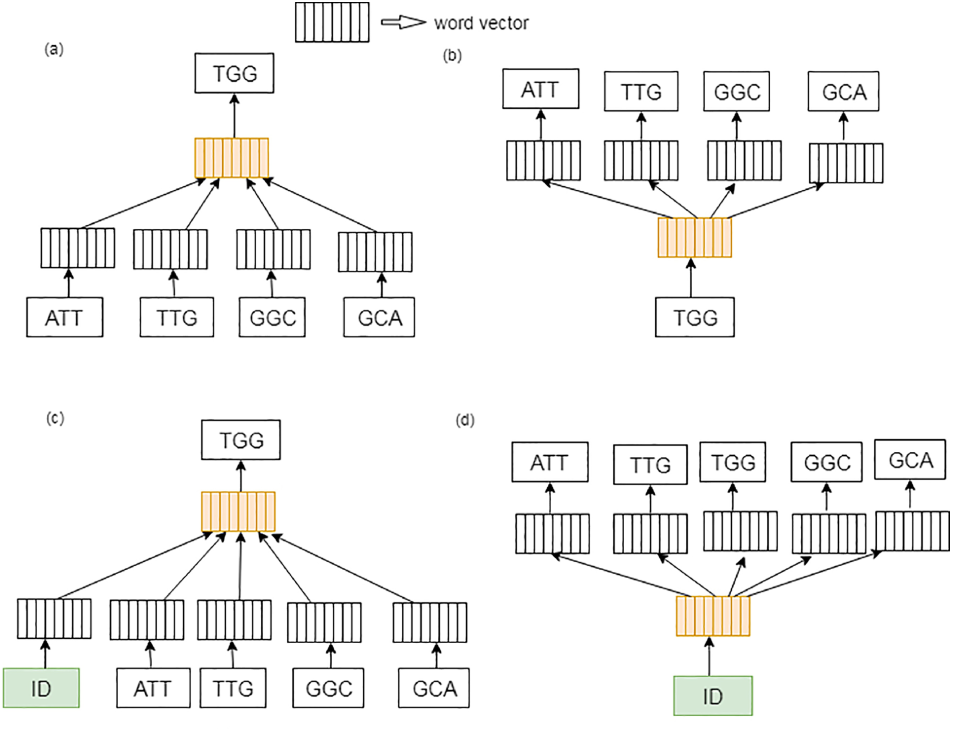
Architecture of word2vec and doc2vec models. (a) CBOW for word2vec (b) skip-gram for word2vec (c) DM for doc2vec and (d) DBOW for doc2vec.

#### 2.3.3. SpliceVec feature space construction

We generated both SpliceVec-g and SpliceVec-sp for each of 587,778 splice junctions, consisting of 50% true and 50% decoy splice junctions. There are several hyper-parameters for both the word2vec and doc2vec models based on which SpliceVec was generated and each of these hyper-parameters needs to be tuned as per the underlying task to be accomplished. We generated several sets of SpliceVec by variations and different combinations of the hyper-parameters mentioned in Table 1. We evaluated the embeddings on the classification task of splice junctions. We partitioned 30% of our dataset as test data. Out of the remaining 70%, 20% was used as validation set for tuning the hyper-parameters of MLP for classification. We trained the MLP with the training data in a Tensorflow [30] implementation. Based on the performance of our classifier, we have fixed the hyper-parameters as given in Table 1. Default values of respective models were considered for other hyper-parameters not mentioned in the table.

**Table 1:**
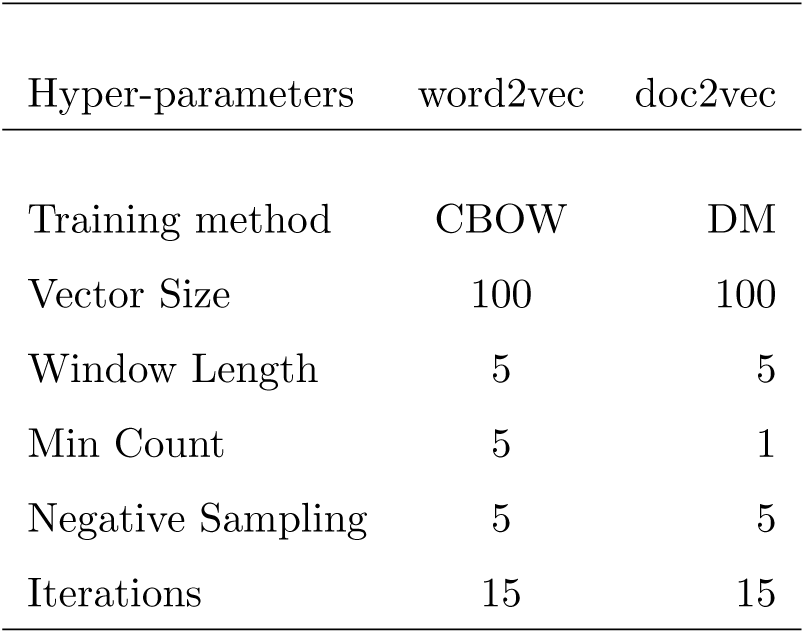
Optimal hyper-parameters for word2vec and doc2vec training models.

| Hyper-parameters | word2vec | doc2vec |
| --- | --- | --- |
| Training method | CBOW | DM |
| Vector Size | 100 | 100 |
| Window Length | 5 | 5 |
| Min Count | 5 | 1 |
| Negative Sampling | 5 | 5 |
| Iterations | 15 | 15 |

### 2.4. Classification of SpliceVec by MLP

We used an MLP with one hidden layer as the classifier for splice junction classification. We take each n-dimensional SpliceVec representing one splice junction as the input data and send the weighted input to each node of the hidden layer. Each node *j* of the hidden layer receives the signal (*x*_*i*_) from each node *i* in input layer multiplied with a weight (*w*_*ji*_). The effective signal *S*_*j*_ of a node *j* is:

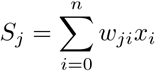

where *n* is the number of nodes in input layer and *x*_0_ is the bias. The signal, in hidden layer, undergoes the activation function. We used rectified linear units (ReLU) as the activation function. ReLU sets any negative input signal (*x*) to zero by the following function:

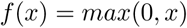

In the output layer, weighted sum of outputs from hidden layer undergoes the softmax activation function that gives the class probabilities of the input. There are two nodes in the output layer corresponding to true and false splice junctions. The predicted output is compared to the expected output using cross entropy as the loss function. The cross entropy loss can be defined as follows:

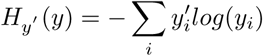

where *y* is the predicted output and *y′* is the expected output. We used Adam Optimizer [31] to minimize the loss by updating weights at each layer.

We compared classification performance of SpliceVec-MLP with existing state-of-the-art approaches for splice site prediction, named SpliceMachine [11] and DeepSplice. SpliceMachine is based on linear support vector machines (LSVM) whereas DeepSplice is based on CNN. We tuned hyper-parameters for DeepSplice, SpliceMachine and SpliceVec-MLP by partitioning training data into train, test and validation set as explained in the previous subsection. The optimal hyper-parameters are given in Table 2. All the experiments were carried out on a 3.20 GHz Quad-Core Intel Core i5 with 8GB memory. We evaluated the performance of the classifier based on precision, recall, accuracy, and F1 score.

**Table 2:** Optimal hyper-parameters of the classi_ers.

| DeepSplice |  | SpliceMachine |  |
| --- | --- | --- | --- |
| Batch size | 160 | Context size (donor) | 20, 60 |
| Dropout keep_prob | 0.8 | Context size (acceptor) | 20, 20 |
| Learning rate | 0.001 | C parameter for LSVM | 0.015625 |
| SpliceVec-MLP |  |  |  |
| Batch size | 128 |  |  |
| Hidden nodes | 1024 |  |  |
| Learning rate | 0.001 |  |  |

## 3. Results

### 3.1. Qualitative analysis of SpliceVec

To analyze the quality of embeddings, we plotted both SpliceVec-g and SpliceVec-sp by projecting it from a 100-dimensional feature space to a 2D space using t-SNE. As a comparison, we plotted an equal number of randomly generated 100-dimensional vectors that are randomly assigned as true or false splice junctions. The 2D embeddings clearly show that both SpliceVec-g and SpliceVec-sp form clusters which suggest that they capture the features specific to true and false splice junctions (Figure 3).

**Figure 3:**
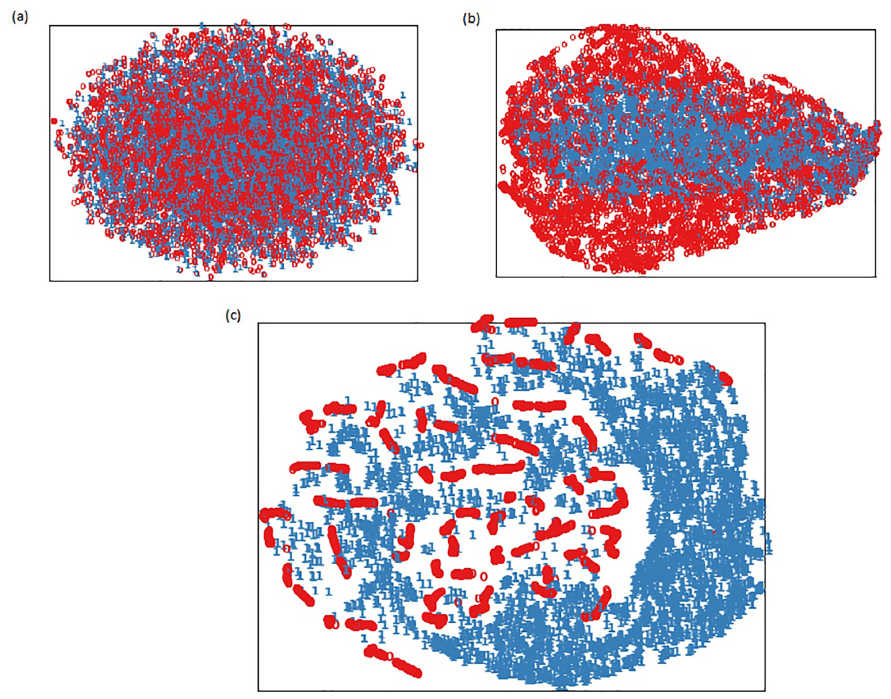
t-SNE plots for different embeddings. (a) Random embedding (b) SpliceVec-g (c) SpliceVec-sp. Each point represents a splice junction. Points in red represent false splice junctions whereas points in blue represent true splice junctions.

We see that SpliceVec-sp displays better partitioning of true and false junctions compared to SpliceVec-g. This is because SpliceVec-sp, based on doc2vec model, captures the order in which words appear in the sequence, thus generating more meaningful feature vector. In Figure 3, points in red represent false splice junctions whereas points in blue represent true splice junctions.

### 3.2. Optimal sequence length for SpliceVec

We varied the length of flanking region at the donor and acceptor splice sites to find the optimal length of sequence that produces the best results in classification. These flanking regions were considered in order to capture the splicing signals present in vicinity of the intron boundary. For each splice junction, we first extracted only 10 nt from upstream and downstream sequence of both donor and acceptor splice sites, thus yielding a sequence length of 40 nt. We increased the length of flanking regions upto 40 nt, in an interval of 10 nt. We also extracted entire intron sequence along with 10 nt upstream and 10 nt downstream of donor and acceptor splice sites respectively. We have taken the entire intron sequence as part of the input because there are evidences of intronic sequences, more than 150 bp long, being conserved around the alternative exons [32, 33], thus making intronic elements more important in regulating AS. In this case also, we increased the length of flanking region from 10 nt upto 40 nt in an interval of 10 nt.

Table 3 demonstrates the results obtained on varying the sequence length. We indeed observe that the improvement in performance of classifier is significantly more when we consider the entire intronic sequence with 10 nt flanking region compared to increasing the length of flanking upstream and downstream regions at both the donor and acceptor splice sites. On further increasing the length of flanking region, in the case where full intron was considered, we observe that performance degrades for SpliceVec-g. This indicates that irrelevant features are being captured as the length of flanking region is increased. On the other hand, SpliceVec-sp is consistent even with increased flanking region. This is because SpliceVec-sp, based on doc2vec model, provides more robust embeddings especially with longer documents and are therefore less sensitive to irrelevant features. We therefore performed our classification task on SpliceVec generated from full intron with 10 nt flanking regions for further analysis. The problem of having very long and variable length introns as input is solved by the fact that each splice junction will be reduced to a 100 dimensional SpliceVec, which is much less than the actual length of intron.

**Table 3:** Performance of splice junction classification using SpliceVec-g and SpliceVec-sp by varying length of junction sequences. We have computed accuracy (Ac), precision (Pr), recall (Re) and F1 score (F1) in percentage as performance measures. The values are average of five simulations.

| Sequence length | SpliceVec-g | SpliceVec-sp |
| --- | --- | --- |
|  | *Ac, Pr, Re, F1(%)* | *Ac, Pr, Re, F1(%)* |
| 10nt flanking | 82.26, 80.61, 85.02, 82.73 | 99.77, 99.80, 99.73, 99.77 |
| 20nt flanking | 81.09, 78.85, 85.01, 81.79 | 99.55, 99.50, 99.61, 99.55 |
| 30nt flanking | 82.68, 80.67, 85.98, 83.22 | 99.52, 99.45, 99.58, 99.52 |
| 40nt flanking | 84.81, 82.87, 87.80, 85.24 | 99.37, 99.33, 99.41, 99.37 |
| 10nt flanking + intron | **98.15, 98.40, 97.89, 98.14** | 99.88, 99.81, 99.95, 99.88 |
| 20nt flanking + intron | 97.73, 97.97, 97.48, 97.72 | 99.94, 99.92, 99.96, 99.94 |
| 30nt flanking + intron | 97.35, 97.86, 96.82, 97.34 | 99.93, 99.92, 99.93, 99.93 |
| 40nt flanking + intron | 97.04, 97.30, 96.77, 97.03 | **99.97, 99.97, 99.97, 99.97** |

### 3.3. Improved classification of splice junctions

We have used SpliceVec as feature in MLP classifier having one hidden layer. Table 4 shows the performance of SpliceVec-g and SpliceVec-sp obtained from the optimal sequence length (10nt flanking + intron) reported in previous sub-section. For SpliceVec-g, our classifier gives an accuracy 2.42-17.13% more than that of the alternative approaches whereas for SpliceVec-sp, our classifier out-performs the counterparts by 4.15-18.86%. SpliceMachine predicts a splice site (either donor or acceptor) given a sequence. Since our approach predicts a junction pair (both acceptor and donor), we therefore calculated the performance of SpliceMachine for junction pair by considering a junction pair as true if both the donor and acceptor sites were predicted as true.

**Table 4:** Performance comparison of different models for splice junction prediction. Performance of SpliceVec is obtained by considering optimal sequence length of 10nt flanking region including complete intronic sequence.

| Model | Performance measures |
| --- | --- |
| | $Ac, Pr, Re, F1(\%)$ |
| SpliceMachine (junction pair) | 81.02, 51.59, 85.41, 64.33 |
| DeepSplice | 95.73, 95.87, 95.60, 95.74 |
| SpliceVec-g - MLP | 98.15, 98.40, 97.89, 98.14 |
| SpliceVec-sp - MLP | 99.88, 99.81, 99.95, 99.88 |

We were also curious to access the performance of SpliceVec for prediction of only donor or acceptor sites. For this, we generated SpliceVec for the consensus dimer (GT for donor and AG for acceptor respectively) along with 10 nt upstream and downstream sequence. It was then classified with SpliceVec-MLP. The results obtained for both SpliceVec-g and SpliceVec-sp for classification of donor and acceptor sites and its comparison with SpliceMachine is shown in Table 5.

**Table 5:** Performance comparison of SpliceVec-MLP with SpliceMachine for prediction of donor and acceptor sites individually.

| Model | Performance measures |
| --- | --- |
| | $Ac, Pr, Re, F1(\%)$ |
| SpliceMachine (donor) | 92.65, 91.36, 94.20, 92.75 |
| SpliceVec-g - MLP (donor) | 84.53, 82.92, 87.02, 84.90 |
| SpliceVec-sp - MLP (donor) | 99.86, 99.82, 99.90, 99.86 |
| SpliceMachine (acceptor) | 87.01, 85.57, 89.04, 87.27 |
| SpliceVec-g - MLP (acceptor) | 80.16, 78.61, 82.89, 80.69 |
| SpliceVec-sp - MLP (acceptor) | 99.84, 99.79, 99.89, 99.84 |

We observe that SpliceVec-sp performs better than SpliceVec-g. This is because SpliceVec-g was generated by computing average of vector representations of all the 3-mers in the sequence. Information regarding the order of the 3-mers was not captured in that case. Whereas, SpliceVec-sp captures the ordering information better because it generates the vector representation of a 3-mer based on its neighboring 3-mers in the sequence.

### 3.4. Robust classification of SpliceVec-MLP

We tested the robustness of SpliceVec-MLP by analyzing its performance with reduced training examples as well as imbalanced training examples. We have varied the number of input samples for training the classifier and observed that SpliceVec-MLP maintained an accuracy more than 98% upto 50% reduction of training samples for SpliceVec-g, whereas for SpliceVec-sp, 99% accuracy was maintained upto 60% reduction of the training samples. On the other hand, the accuracy of DeepSplice reduced by about 6% (Table 6). Observing the accuracy of SpliceVec models with reduced dataset, we can conclude that SpliceVec captured more meaningful features compared to DeepSplice.

**Table 6:** Classification by reducing training dataset: Performance comparison of DeepSplice and SpliceVec.

| Train data (%) | DeepSplice | SpliceVec-g | SpliceVec-sp |
| --- | --- | --- | --- |
|  | *Ac, Pr, Re, F1*(%) | *Ac, Pr, Re, F1*(%) | *Ac, Pr, Re, F1*(%) |
| 100 | 95.73, 95.87, 95.60, 95.74 | 98.15, 98.40, 97.89, 98.04 | 99.88, 99.81, 99.95, 99.88 |
| 70 | 93.90, 94.80, 92.90, 93.84 | 98.05, 97.77, 98.28, 98.05 | 99.94, 99.91, 99.98, 99.94 |
| 60 | 93.40, 93.65, 93.11, 93.38 | 98.02, 98.28, 97.74, 98.02 | 99.67, 99.96, 99.38, 99.67 |
| 50 | 90.26, 90.24, 90.27, 90.26 | 98.05, 98.43, 97.65, 98.04 | 99.62, 99.28, 99.97, 99.63 |
| 40 | 89.79, 88.95, 90.87, 89.90 | 97.98, 97.42, 98.57, 97.99 | 99.88, 99.93, 99.83, 99.88 |

To analyze the performance of SpliceVec on imbalanced classes, we varied the ratio of negative to positive samples from 5 to 17 in an interval of 3. This analysis is particularly important because in real scenario the number of negative samples is much more than that of positive samples in a genome. Figure 4 shows that the performance of both SpliceVec-g and SpliceVec-sp is significantly more consistent with increasing ratio of negative samples as compared to DeepSplice.

**Figure 4:**
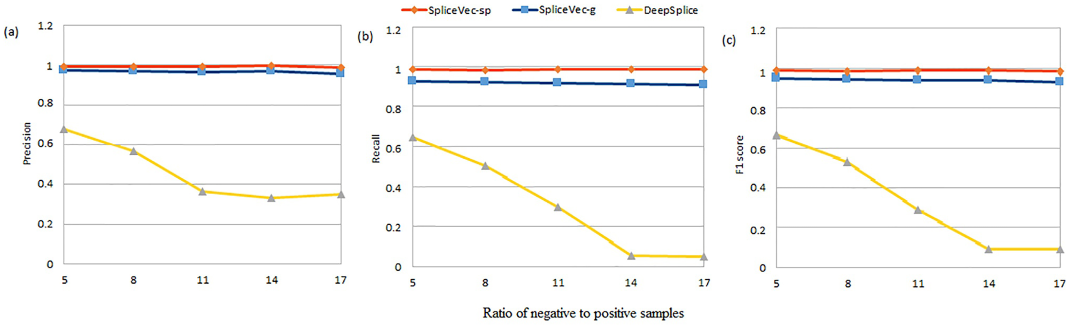
Performance of SpliceVec with increasing ratio of negative samples to positive samples. (a) Precision (b) Recall (c) F1-score of SpliceVec-sp, SpliceVec-g and DeepSplice on varying ratio of negative samples from 5 to 17. Data here is positive and negative samples from chromosome 20 of GENCODE dataset.

### 3.5. Prediction performance

The proposed models performed better than state-of-the-art model in detecting non-canonical true splice junctions. SpliceVec-g identified upto 192 (84.95%) out of 226 non-canonical true splice junctions present in our test dataset. SpliceVec-sp identified all the 226 (100%) non-canonical splice junctions. The capability of the classifier to detect non-canonical splice junctions significantly improved on including complete intron sequence for generating SpliceVec.

Observing 100% accuracy in detecting non-canonical splice junctions, we further modified our decoy splice junction samples by including non-consensus donor and acceptor dimers (dimers other than GT-AG pair) in it. In order to include non-consensus dimer-pairs, we considered two common classes of exceptions to consensus splice site sequences reported in [34]. We considered 0.5% AT-AC dimer-pairs and 0.5% GC-AG dimer-pairs based on their reported frequency of occurrence. In this scenario, SpliceVec-g identified 188 (83.18%) out of 226 non-canonical true splice junctions whereas SpliceVec-sp identified all the 226 (100%) junctions in the worst case out of 5 simulations (considering 10 nt flanking region + intron). Performance of SpliceVec-MLP indicates that the feature representation by SpliceVec is invariant to canonical and non-canonical splice junctions.

We also observed that the most common correctly identified dinucleotide sequence at the donor site of non-canonical splice junction was AT whereas that at the acceptor site was AC, constituting about 43% of the non-canonical splice junctions. This is in consistency with experimentally studied annotation which identifies AT-AC introns as one of the most important classes of exception to splice site consensus [34]. Table 7 shows the prediction of true splice junctions in the worst case out of 5 simulations on varying the sequence length for both SpliceVec-g and SpliceVec-sp. Out of the 87,927 canonical and 226 non-canonical true splice junctions, DeepSplice identified 84,170 (95.73%) canonical and 97 (42.92%) non-canonical splice junctions. Thus, SpliceVec-MLP showed 2.12% (4.23%) higher performance than DeepSplice in terms of identification of canonical splice junctions using SpliceVec-g (SpliceVec-sp). In terms of identification of non-canonical splice junctions, there was a 42.03% (57.08%) improvement in performance by SpliceVec-g (SpliceVec-sp) compared to DeepSplice.

**Table 7:** Correctly predicted canonical (*Pred*_*can*_) and non-canonical (*Pred*_*non-can*_) positive splice junctions, out of 87,927 canonical and 226 non-canonical junctions, using MLP and both SpliceVec-g and SpliceVec-sp by varying the length of junction sequence.

| Model | SpliceVec-g |  | SpliceVec-sp |  |
| --- | --- | --- | --- | --- |
| | $Pred_{can}$ | $Pred_{non-can}$ | $Pred_{can}$ | $Pred_{non-can}$ |
| 10nt flanking | 72,446 | 51 | 87,635 | 222 |
| 20nt flanking | 74,172 | 84 | 87,375 | 223 |
| 30nt flanking | 75,222 | 127 | 87,367 | 222 |
| 40nt flanking | 77,794 | 141 | 87,098 | 223 |
| 10nt flanking + intron | 86,037 | 190 | 87,854 | 226 |
| 20nt flanking + intron | 85,656 | 192 | 87,884 | 226 |
| 30nt flanking + intron | 84,814 | 191 | 87,888 | 226 |
| 40nt flanking + intron | 84,190 | 186 | 87,899 | 225 |

We compared the time taken for classifying the test samples. SpliceVec-MLP classified 72,268 test samples per second of CPU time in the worst case out of 5 simulations. DeepSplice, on the other hand, classified 5,585 test samples per second. SpliceVec-MLP was therefore 12.94 times faster than DeepSplice. With SpliceVec-sp, the classifier could identify all the splice junctions belonging to some of the important genes like *TSLP* and *BATF2* that are known to be involved in several diseases. *TSLP* causes diseases like atopic dermatitis, eosinophilic esophagitis, and allergic rhinitis [35]. *BATF2* has been known to be associated with cancer and some allergic diseases [36].

## 4. Conclusions

We proposed a novel approach of feature representation for splice junctions, named SpliceVec, which is a vector embedding of the junction in an n-dimensional feature space that captures the splicing signals residing in vicinity of the splice junctions. We classified SpliceVec with an MLP having single hidden layer and showed that it outperforms the current state-of-the-art splice junction prediction model even with a significantly reduced training dataset. The model performs consistently even with imbalance in training samples. This representation also performs outstanding for prediction of non-canonical true splice junctions. We have also seen that this classification is computationally times faster than the state-of-the-art model thus making this model scalable to the availability of annotated genome sequences. Although SpliceVec has been trained for human genome, it can easily be trained with user-defined training data or data belonging to any other species.

## Abbreviations

AS: alternative splicing
CNN: convolutional neural network
t-SNE: stochastic neighbor embedding
MLP: multilayer perceptron
nt: nucleotides
bp: base pairs
NLP: natural language processing
CBOW: continuous bag of words
DBOW: distributed bag of words
DM: distributed memory
ReLU: rectified linear units
LSVM: linear support vector machines

